# A new animal model signifying a decisive bridge between innate immunity and the pathognomonic morphological characteristics of Type 1 Diabetes

**DOI:** 10.1101/2021.12.10.472088

**Authors:** Tegehall Angie, Ingvast Sofie, Melhus Åsa, Skog Oskar, Korsgren Olle

**Affiliations:** Department of Immunology, Genetics and pathology, Rudbeck Laboratory, Uppsala University, 751 85 Uppsala, Sweden; Department of Medical Sciences, Section of Clinical Microbiology, Uppsala University, Uppsala, Sweden

## Abstract

Available animal models for Type 1 Diabetes (T1D) show limited similarities with the human disease and have no predictive value in screening for effective intervention therapies. Heat-inactivated bacteria instilled in the ductal compartment of the pancreas in healthy rats rapidly cause periductal inflammation and accumulation of mainly granulocytes and monocytes in the exocrine pancreas and in the peri-islet area mimicking the acute pancreatic inflammation in subjects with recent onset T1D. After three weeks, the triggered exocrine inflammation had vanished and pancreases showed normal morphology. However, a distinct accumulation of both CD4+ and CD8+ T cells within and adjacent to affected islets was found in one third of the rats, mimicking the pathognomonic insulitis in T1D. As in human T1D, the insulitis affected a fraction of all islets and was observed only in certain lobes of the pancreases. The presented animal model for T1D will allow detailed mechanistic studies to unravel a previously unknown interplay between bacteria-activated innate immunity and an acquired cellular immunity forming the immunopathological events described in humans at different stages of T1D.

**Summary Statement:** The results presented signify a previously unknown decisive bridge between innate immunity and formation of the pathognomonic immunopathological events described in subjects with recent onset T1D.

## Introduction

The autoimmune barrier in Type 1 Diabetes (T1D) is only vaguely understood. However, an immunopathological hallmark in the pancreas of patients with T1D is the accumulation of immune cells within and around affected islets, i.e., insulitis (Campbell-Thompson et al., 2016a, Reddy et al., 2015). Notably, only few T cells infiltrate the islets (In’t Veld, 2014, Krogvold et al., 2016, Reddy et al., 2015). In fact, an extensive study using multiplex immunofluorescent staining of 35 simultaneous biomarkers with a spatial resolution of 1 μm demonstrated that immune and islet cells essentially remain isolated from each other even in patients with recent onset T1D (Damond et al., 2019). Phenotypically, CD8^+^ and CD4^+^ T cells dominate the insulitis, followed by monocytes/macrophages. B cells are less frequent, and regulatory T cells, NK cells and plasma cells are only rarely found (Willcox et al., 2009). Also, the exocrine pancreas is affected, as evidenced for example by a markedly smaller volume and presence of multifocal T-cell infiltrates in acinar regions.

The proportion of islets with insulitis has been inversely associated with disease duration in some (Campbell-Thompson et al., 2016a), but not other studies (Reddy et al., 2015). Insulitis seems not related to age at onset, number of autoantibodies, or HLA genotype (Campbell-Thompson et al., 2016a). The presence of remaining insulin-positive cells in islets with insulitis several years, or even decades, after diagnosis of T1D is intriguing and argue against a classical T-cell mediated cytotoxic elimination of the insulin-producing cells. On the contrary, our current knowledge of T1D suggests a mild and slowly progressing disease.

A substantial proportion the CD8+ T cells in the insulitic lesions from subjects with recent onset T1D constitute tissue resident memory T cells (T_RM_ cells) (Kuric et al., 2017). T_RM_ cells were present in all T1D subjects examined and represented about 40% of the total number of CD8^+^ T cells per inflamed islet, a proportion of T_RM_ cells similar to that previously described in skin lesions of psoriasis (Cheuk et al., 2014). Also, T and B cell gene expression pattern in infiltrated islets argue against that the T cells found in the insulitic lesions constitute conventional cytotoxic T CD8+ cells (Krogvold et al., 2016). T_RM_ cells constitute a subset of memory T cells that persist for years at the site of a previous infection without persistence of antigen stimulation and provide rapid immune protection against re-infection via the same entry port (Clark, 2015). The substantial proportion of T_RM_ cells in islets of recent onset T1D subjects support the hypothesis of infectious agents in the development of T1D (Skog et al., 2013).

Progress in medical research often depend on the availability of validated small animal models, especially so in disease-oriented research aiming to develop effective intervention therapies, including T1D. Lack of predictive and validated animal models has indeed caused the pharmaceutical industry to abandon entire disease areas (Pound, 2020).

The most common animal models in T1D research are the NOD mouse and the BB rat. Unfortunately, both models show limited similarities with human T1D and have little predictive value in screening for effective intervention therapies (Roep and Atkinson, 2004). Major dissimilarities between these animal models and human T1D include 1) the general dysregulation of their immune system causing disease progression in several organs besides the pancreas, 2) the gender and strain-dependency, 3) the dependency of specific husbandry conditions, and 4) major histopathologic differences in morphology and immune mediated destruction of the beta cells when compared to that in humans.

The hallmark of T1D is a slowly (years) progressing decline in endogenous insulin secretion resulting in an increased blood glucose. In the search for an animal model for T1D, high blood glucose has often been the primary criterion. However, rodents, in contrast to humans, have a remarkable capacity to regenerate both the exocrine and endocrine pancreas, e.g. three weeks after 90% pancreatectomy in the rat both the exocrine and endocrine volumes are almost restored (Brockenbrough et al., 1988). This difference in regenerative capacity should be considered together with the remitting and relapsing disease process over many years and the patchy distribution of affected areas of the pancreas in humans affected by T1D (Gepts, 1965), i.e. at each point of time, some lobes of the pancreas are affected whereas others remain apparently intact. This patchy affection of the pancreas is one of the most characteristic immunopathological findings in T1D and shows resemblances with several other immune mediated diseases, including psoriasis, alopecia and multiple sclerosis. A remitting/relapsing and patchy disease process in a species with significant regenerative capacity would trigger a compensatory formation of new islets from unaffected areas of the pancreas thereby preventing development of hyperglycemia. In fact, in NOD mice or BB rats all beta cells in the entire pancreas are destroyed over a short period of time, an immune-process vastly different from that in human T1D (Alanentalo et al., 2010). Therefore, we should refrain from including overt diabetes as the main criterion for clinically relevant rodent models of T1D.

We have previously reported that injection of heat-inactivated human pathogens in the ductal compartment of the pancreas in healthy rats of several different strains causes periductal inflammation and accumulation of immune cells, mainly granulocytes and monocytes, in certain lobes of the exocrine pancreas and in the peri-islet area (Korsgren et al., 2012). Small bleedings or large dilatations of the capillaries were frequently found within the islets and several beta cells showed severe hydropic degeneration, i.e., swollen cytoplasm, but with preserved nuclei. These findings show marked similarities with those observed in the pancreases of patients dying at onset of T1D. However, inflammation of the pancreas in humans examined weeks or months after diagnosis of T1D is substantially reduced and mainly consists of discreet insulitis, mainly consisting of T cells, in only few islets (Campbell-Thompson et al., 2013, Krogvold et al., 2016).

The present investigation was undertaken in order to study how the acute inflammation triggered by instillation of heat-inactivated bacteria in the ductal compartment develops over a period of several weeks in the rat and how observed findings relate to T1D in humans with an aim to further validate the model for T1D research.

## RESULTS

### Bacterial Challenge and Animal Well-being

All animals tolerated the surgical procedure well. No macroscopic changes were observed in any abdominal organ after 3 or 6 weeks after the bacterial challenge. IVGTT revealed no significant differences in peak glucose values and subsequent glucose disposal 3 or 6 weeks after the bacterial challenge when compared with control rats (p>0.05, Fig. 1).

**Figure 1.**
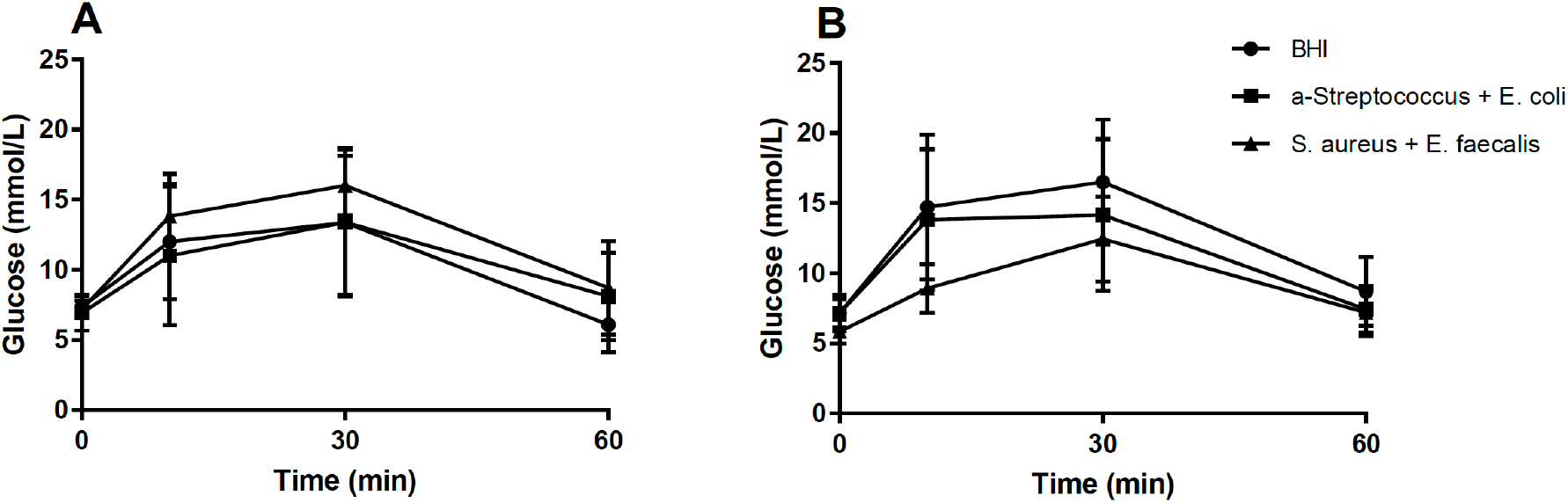
Intravenous glucose tolerance test revealed no significant differences in peak glucose values and subsequent glucose disposal 3 weeks **(A)** or 6 weeks **(B)** after bacterial challenge with *S. aureus* and *E. faecalis* (filled squares) or *E. coli* and α-hemolytic streptococcus (filled triangles) when compared with rats instilled with brain heart infusion broth alone (filled circles; p>0.05).

### Early Innate Antibacterial Responses

Pancreatic sections from a human organ donor with acute onset T1D showed a >2-fold overexpression of 18 antibacterial genes compared to non-diabetic donors (P<0.05, fdr<10%) (Fig. 2A). Bacterial translocation in the rat induced after 4 h a >2-fold overexpression of 22 genes and underexpression of one gene (ccl5/rantes) related to antibacterial response (P<0.05, fdr<10%) (Fig. 2B). Eight of these where homologues to genes that were significantly induced by bacterial translocation in the human.

**Figure 2.**
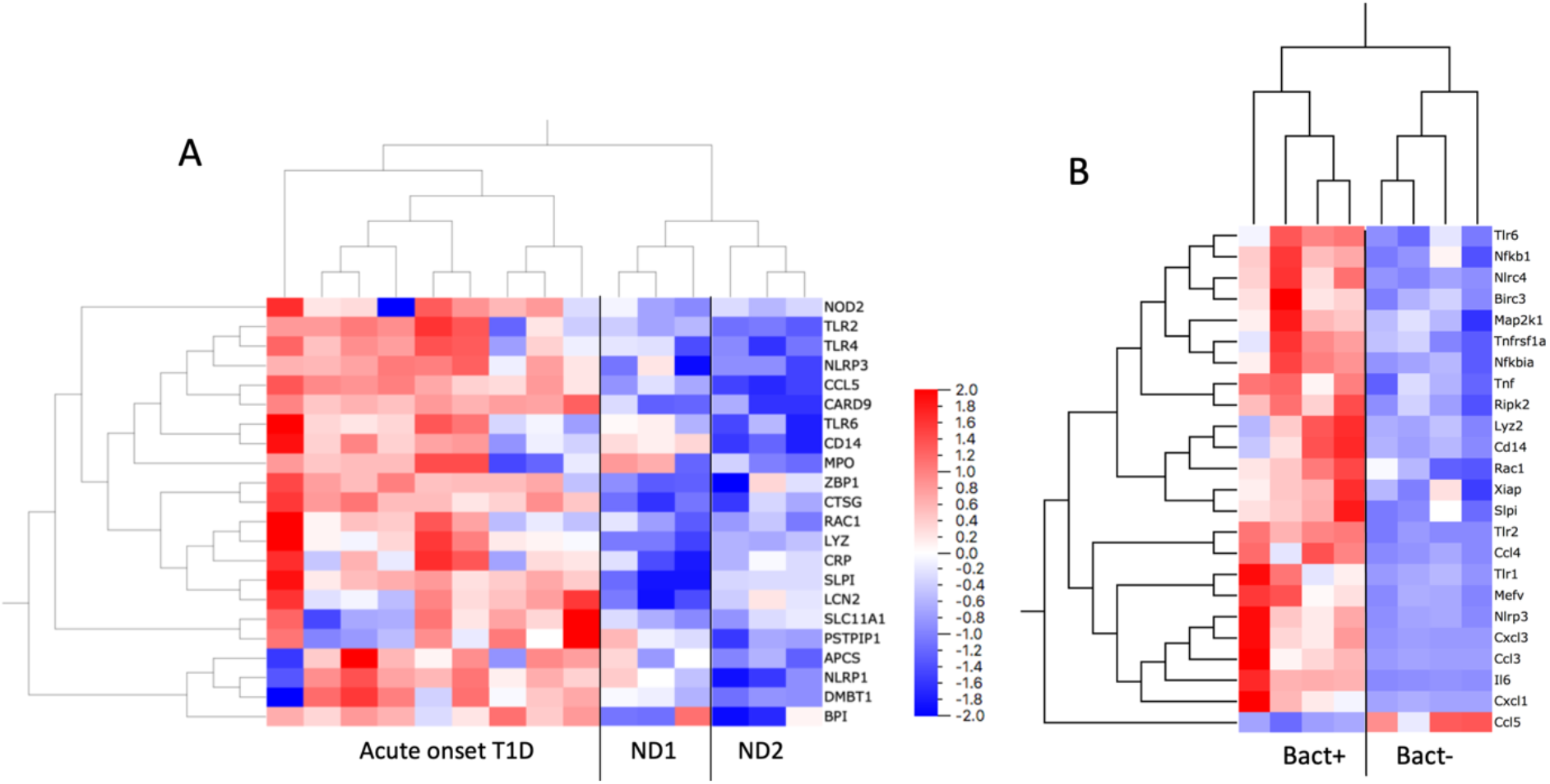
**A)** Expression of 84 human genes related to innate antibacterial response was analyzed with a qPCR array. Pancreatic RNA extracted from an organ donor that died at acute onset of T1D showed a >2-fold overexpression of 18 antibacterial genes compared to non-diabetic donors (ND1 & ND2) (P<0.05, fdr<10%). (**B)** Expression of 84 rat genes related to innate antibacterial response were analyzed with a qPCR array. RNA was extracted from pancreatic rat tissue with (bact+) or without (bact-) instilled bacteria. Bacterial translocation induced a >2-fold overexpression of 22 genes and under expression of one gene (ccl5/rantes) related to antibacterial response (P<0.05, fdr<10%). Eight of these were homologues to genes that were significantly induced by bacterial translocation in the human tissue samples.

### Immune-cell Infiltration and Insulitis

#### A human organ donor that died at onset of type 1 diabetes

A detailed morphological description of the inflammation in the pancreas of the 29-year-old organ donor that died at onset of T1D has been reported previously (Korsgren et al., 2012). This donor fulfilled the consensus definition of insulitis (Campbell-Thompson et al., 2013) and clusters of immune cells were frequently observed close to the islets (insulitis) (Fig. 3A). Immunohistochemical staining for CD3 revealed that most cells in the insulitis were T cells. Of these, a majority was CD8^+^, but several CD4+ cells were also observed (Korsgren et al., 2012).

**Figure 3.**
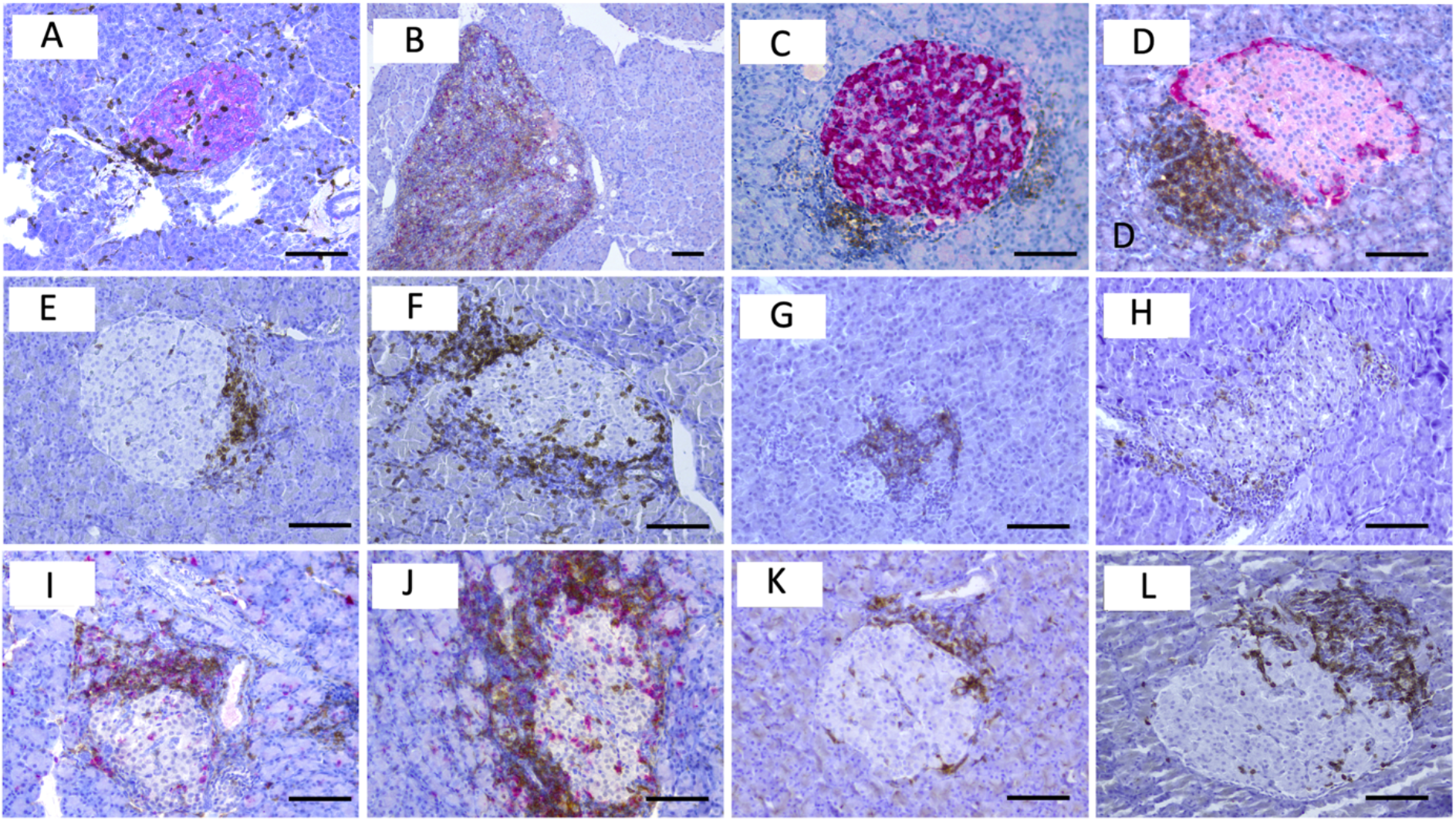
Pancreatic tissue from a 29-year old organ donor who died at onset of type 1 diabetes **(A)** or from rats three weeks after installation of *E. faecalis* in the pancreatic duct **(B-L)** stained for islet hormones and immune cell markers. **A)** An islet with insulitis; CD45 brown, synaptophysin red. **B)** focal area of dense accumulation of immune cells resembling the organization of lymphatic tissue; CD4 brown, CD8 red. **C-L)** Islets with varying degrees of insulitis; **C)** insulin red, CD43 brown. **D)** glucagon red, CD43 brown. **E-F)** CD3 brown, **G)** CD20 brown. **H)** CD68 brown. **I-J)** CD4 brown, CD8 red. **K-L)** CD103 brown. Scale bars represents 100 μm.

#### Rats three weeks after bacterial challenge

The general pancreatic inflammation seen in rats acutely after bacteria translocation was not different when compared to our previous report (Korsgren et al., 2012). This acute inflammation in response to the bacterial challenge was no longer observed three weeks after translocation. Seven of the eight control animals had normal pancreatic architecture with presence of only occasional immune cells. However, some of the rats showed increased occurrence of fibrosis in the head of the pancreas and one control animal had signs of a small bleeding in the head region of the pancreas with presence of small numbers of CD68+, CD20+, CD3+ and CD4+ and CD3 and CD4+ immune cells. The pancreatic tail region of this animal had very few immune cells and no signs of bleedings.

Five of the eight animals treated with ductal instillation of a bacterial mixture showed normal pancreatic architecture with presence of only occasional immune cells. However, in three of the rats injected with *E. faecalis*, some pancreatic lobes in both the head and tail regions contained occasional focal areas of dense accumulations of immune cells (Fig. 3B), resembling the organization of lymphatic tissue with presence of large numbers of T cells (CD3+, CD8+ and CD4+) and B cells (CD20+). These areas of dense lymphatic tissues seemed randomly distributed in the exocrine pancreas.

The most noticeable finding in these three rats was presence of insulitis in some lobes of both the head and tail regions of the pancreases. In the affected regions, 20-80% of islets showed insulitis (Fig. 3C-L). Islet architecture remained normal with presence of insulin-positive cells preferentially in the center and glucagon-positive cells preferentially in the periphery of the islets (Fig 3C-D). Phenotypically, a majority of the immune cells in the insulitic lesions were T cells (CD3^+^) (Fig. 3E-F), but also B cells (CD20^+^) (Fig. 3G) and macrophages (CD68^+^) (Fig. 3H) were common. Both CD8+ and CD4+ cells were frequent among the T cells in the insulitic lesions (Fig. 3I-J) and many displayed the tissue residence marker CD103 (Fig. 3K-L).

#### Rats six weeks after bacterial challenge

All animals had normal pancreatic architecture with presence of only occasional immune cells. No rat showed insulitis. However, half of the rats challenged with heat-inactivated bacteria in the ductal system showed increased occurrence of fibrosis in the head of the pancreas.

## DISCUSSION

In the present study, a new rat model in T1D was validated against the classical morphological hallmarks of human T1D. The intense cellular innate inflammation triggered acutely after instillation of heat-inactivated bacteria in the ductal compartment (Korsgren et al., 2012) was seemingly resolved after 3 weeks. However, in about one third of the animals examined after 3 weeks, a distinct accumulation of both CD4+ and CD8+ T cells was found within and adjacent to some of the islets, fulfilling the consensus definition of insulitis in humans (Campbell-Thompson et al., 2013). As in humans with recent onset T1D, the insulitis affected a minority of islets and was observed only in certain lobes of the pancreases examined.

No deterioration of glucose metabolism was found in rats subjected to an IVGTT. At first this may seem disappointing, however, a rodent model mimicking human T1D should in the initial disease process affect only some islets. It could be speculated that at least two additional circumstances are required for hyperglycemia to develop; 1) repeated lesions that over time that would gradually affect increasingly larger volumes of the pancreas and 2) a species unable to regenerate the pancreas between each insult. Rodents able to restore both endocrine and exocrine pancreas in just a few weeks (Brockenbrough et al., 1988) are therefore not suitable for studying the effects on long-term glucose metabolism. It is also plausible that an animal less resilient to bacterial challenges than the rat, e.g., humans, would show more negative effects on their glucose metabolism.

Instillation of heat-inactivated bacteria induced a self-resolving short-term patchy innate inflammation in the rat. However, about one third of the rats showed induction of acquired immunity as evidenced by persistent insulitis consisting mainly of CD8+ T cells. This type of insulitis is seemingly identical to that found in human subjects examined a few weeks after diagnosis of T1D, e.g. in the DiViD biopsy study (Krogvold et al., 2016, Kuric et al., 2017). Notably, a fraction of the T cells found in the insulitic lesions in the rats expressed the CD103 antigen. The precise role of CD103 on T cells is not fully understood. However, it has been suggested that CD103 is important for conjugation of CD8^+^ T cell to E-cadherin-expressing epithelial cells, thereby facilitating their destruction upon virus and bacterial infections (Clark, 2015). Recently, we reported on the presence of significant number of tissue resident memory T cells (CD8+, CD69+, CD103+) in the insulitic lesions in humans with recent onset T1D (Kuric et al., 2017) similar to the herein reported expression of CD103 on T cells in the insulitis in rats after bacterial instillation. Notably, also the areas of dense lymphatic tissues, seemingly randomly distributed in the exocrine pancreas in the rat model mimic the compartmentalized tertiary lymphoid organs recently reported in human subjects with short T1D disease duration (Smeets et al., 2021, Korpos et al., 2021).

Importantly, half of the rats examined 6 weeks after installation of heat-inactivated bacteria in the ductal compartment showed increased fibrosis in some lobes of the exocrine pancreas. The involvement of the exocrine pancreas in patients with T1D is underappreciated; several studies have reported the appearance of autoantibodies against exocrine cells prior to the onset of T1D (Hardt et al., 2008, Panicot et al., 1999, Taniguchi et al., 2001, Taniguchi et al., 2003), with high predictive value for the future development of T1D. Focal lesions of acute pancreatitis and an accumulation of leukocytes, often around the ducts, has frequently been reported in a majority of subjects with recent-onset (T1D) (Gepts, 1965, Foulis et al., 1986), and most T1D patients display extensive periductal and interlobular fibrosis, i.e., the end stage of inflammation (Meier et al., 2005). Also, a substantial reduction (≈30%) in pancreatic volume is present in newly diagnosed subjects (Gaglia et al., 2011, Williams et al., 2007, Williams et al., 2012), as well as mild-to-moderate exocrine pancreatic insufficiency (Creutzfeldt et al., 2005). Reduced pancreatic weight has also been demonstrated in pancreases from organ donors with “pre-diabetes” (presence of islet auto-antibodies) (Campbell-Thompson et al., 2016b, Campbell-Thompson et al., 2012).

A role for bacteria in the development of T1D is emerging (Korsgren et al., 2012, Rouxel et al., 2017, Knip and Siljander, 2016, Abdellatif et al., 2019). Notably, both the type and the intensity of the proinflammatory responses induced in isolated human islets depend on the specific strain of bacteria applied (Korsgren et al., 2012, Abdellatif et al., 2019). Although the present study includes only a few bacterial species, only rats instilled with a bacterial challenge including heat-inactivated *E. faecalis* developed the morphological characteristics of T1D. Expression of genes related to anti-bacterial response was upregulated in the pancreas of the subject with recent onset T1D as well as in rats exposed to heat-inactivated bacteria relative to that observed in controls (Fig 2). It is therefore postulated that the observed findings result as an interplay between an external inflammatory trigger (the heat-inactivated bacteria) and a proinflammatory response induced in affected islets (release of cytokines and chemokines).

Trafficking and activation of leukocytes is controlled by chemokines and cytokines produced by parenchymal cells in response to inflammation. The HLA genotypes conferring an increased risk for T1D are linked to increased innate responses to bacterial infections (Kotb et al., 2002) further underlying the importance of the interplay between innate and acquired immune responses in T1D. Several proteins with this powerful immunoregulatory capacity are produced by human islet cells, e.g. CCL2 (MCP-1), CCL5 (RANTES), CCL3 (macrophage inflammatory protein 1-alpha (MIP-1-alpha)), CXCL2 (MIP-2), CXCL9 (monokine induced by gamma interferon (MIG)), CXCL10 (Interferon gamma-induced protein 10 (IP-10)), CXCL11 (Interferon-inducible T-cell alpha chemoattractant (I-TAC)), Macrophage migration inhibitory factor (MIF), IL-1β, IL-6 and IL-8 (Waeber et al., 1997, Johansson et al., 2003, Eizirik et al., 2012). This cascade of cytokines and chemokines released by islet cells in response to inflammation can supposedly explain the constant and puzzling finding, observed already after a few hours, with accumulation of large number of granulocytes and monocytes in the peri-islet area also in lobes not, or only marginally, affected by the inflammation in the exocrine pancreas (Korsgren et al., 2012). At later stages this initial innate islet inflammation is replaced by T and B lymphocytes to form the archetypical insulitis observed in subjects with recent onset T1D (Gepts, 1965). This interplay between the islets and the immune system is illustrated by the distinct insulitis affecting a substantial number of islets also in the splenic part of the pancreas, i.e., parts of the pancreas most distal from the intestine and the injection site of the heat-inactivated bacteria.

Whenever repeated, similar processes would be initiated in additional lobes of the pancreas eventually affecting large volumes of the pancreas. Notably, repeated pancreatic inflammation would likely induce activation of the CD103+ T cells residing in the previously formed insulitic lesions resulting in cytolysis until the total number of beta cells is too low to maintain glucose metabolism. Tentatively, the progression to diabetes is facilitated by the observed periductal fibrosis, the end-stage of periductal inflammation found both in the herein described rat model and in human T1D (Gepts, 1965, Meier et al., 2005), negatively affecting the formation of new islets (Butler et al., 2003, Skog and Korsgren, 2020).

In summary, we present a novel rodent model for the early immunopathological events in T1D. With the data presented here, together with our previous publication (Korsgren et al., 2012), it is demonstrated that this model fulfills the following criteria of being relevant for human T1D: 1) no strain restriction, 2) similar affection of both male and female animals, 3) no dependency of husbandry conditions, 4) dependency on environmental trigger(s), 5) remitting and relapsing disease process, 6) patchy affection of the pancreas, 7) initial innate inflammation of some pancreatic lobes, and 8) subsequent formation of insulitis, a pathognomonic morphological characteristics of T1D. The presented model allows detailed mechanistic studies to unravel the interplay between the innate and acquired immunity in the formation of immunopathological events seemingly identical to those described in humans with recent onset T1D.

## Materials and Methods

### Ethics

All work involving human tissue was conducted according to principles expressed in the Declaration of Helsinki. The consent to use pancreatic tissue from deceased organ donors for research purposes was obtained verbally from the deceased person’s next of kin by the physician in charge or obtained from an online database and fully documented in accordance with Swedish law and regional standard practices. The study was approved by the Regional Ethics Committee in Uppsala, Sweden (Dnr 2009/043, 2009/371, 2015/444). Animal experiments were approved by the Uppsala Laboratory Animal Ethical Committee (permit number C141/15), on the condition that only dead bacteria were used if the observation period was prolonged compared with the initial study (Korsgren et al., 2012).

### Human pancreatic samples

Pancreatic tissue from a 29-year old organ donor that died at onset of T1D (a previously healthy man with B-glucose 46 mmol/L and ketoacidosis at arrival to the emergency room and with a BMI of 24.2 kg/m^2^ and HbA1c 90 mmol/mol, previously described in detail (Korsgren et al., 2012) and two donors without diabetes (a 31 year old woman with BMI 25.4 kg/m^2^ and HbA1c 33 mmol/mol, and a 27 year old man with BMI 26.0 kg/m^2^ and HbA1c 39 mmol/mol), procured within the Nordic Network for Clinical Islet Transplantation, were included in the study. Biopsies were formalin fixed and paraffin-embedded or frozen in liquid nitrogen and stored at −80°C.

### Bacteria

All four strains used in an earlier study in this new model were included (6). They were isolated from patients with invasive infections at the Department of Clinical Microbiology, Uppsala University Hospital, Uppsala, Sweden, and chosen for their documented ability to translocate into pancreas and cause infections in this anatomic region (Flores et al., 2003, Negm et al., 2010, Schmid et al., 1999, Stelzmueller et al., 2007). The bacteria were grown overnight in brain heart infusion (BHI) broth (Becton Dickinson) or Trypticase soy broth BBL with 10% inactivated horse serum and 5% Fildes enrichment BBL at 35°C to a concentration of 10^9^ CFU/mL. After heat-inactivation by boiling for 15 min, the viability was controlled. The dead bacteria were stored at −70°C until used.

### Animals and Operating Procedure

Healthy, male Wistar rats weighing 250 to 300 g (Taconic, Denmark) were used. Before the bacterial challenge, the animals were kept under standard laboratory conditions in accordance with the National Institute of Health principles of laboratory animal care and national laws in Sweden. The rats were housed two by two in plastic cages under a 12:12-h light-dark cycle, and they were given water and food *ad libitum*. At challenge, 200 μl of BHI broth with or without bacteria was instilled as previously described (Korsgren et al., 2012). Animals were subsequently kept under normal conditions for 4 h, 3 or 6 weeks, respectively. At the end of the experiment after 3 or 6 weeks, glucose tolerance was evaluated on some of the rats by an intravenous glucose tolerance test IVGTT under full anesthesia (thiobutabarbital sodium administered 10 minutes before glucose injection (100 mg/kg BW intraperitoneally). Bolus injection of glucose was given within 60 seconds via the tail vein. Blood glucose was measured immediately before and 5, 10, 30, 60, 90 and 120 minutes after glucose injection. Blood glucose was measured with a glucometer (CONTOUR^®^, Bayer, Solna, Sweden), operating within a range of 0.6-33.3 mmol glucose/L.

Animals were subsequently killed by heart puncture and serum, plasma and pancreas were collected. The head and tail of the pancreas were fixed in 4% paraformaldehyde and prepared for paraffin embedding.

### RNA Extraction and qPCR Array

Frozen tissue biopsies from three different parts of the pancreatic body and tail from the organ donor with recent onset T1D and one biopsy from each of the two non-diabetic organ donors were subjected to sectioning and RNA extraction. Twenty consecutive 10 μm sections were placed on glass slides for subsequent IHC (sections 1-2, 7-8, 13-14, & 19-20) or placed in 600 μl buffer RLT (Qiagen) containing 1% 2-mercaptoethanol (Sigma-Aldrich) for extraction of RNA. From each of the five tissue biopsies, sections 3-6, 9-12, and 15-18 were pooled and RNA extracted separately. Frozen tissue from the head and the tail of the pancreas of four rats sacrificed 4 h after the instillation of *Enterococcus faecalis* in the ductal system were subjected to sectioning and RNA extraction following the same protocol as for the human samples.

AllPrep Mini kit (Qiagen) was used for RNA extraction according to the manufacturer’s instructions, including homogenization using QiaShredder columns (Qiagen) and on-column DNase digestion. In the final step, an elution volume of 30 μL was used, giving RNA concentrations ranging from 63 to 217 ng/ μL per sample.

Pathway-specific primer mixes (Rat Antibacterial Response, PBR-148Z, and Human Antibacterial Response, PBH-148Z; Qiagen) were used for preamplification and qPCR arrays (Rat Antibacterial Response, PARN-148ZE, and Human Antibacterial Response, PAHS-148ZC; Qiagen) were used for the expression analysis of 84 genes involved in innate antibacterial responses in human and rat respectively. Genes with a quantification cycle (Cq) value >35 were regarded as non-detected and assigned a Cq of 35 to calculate fold induction.

### Immunohistochemistry

Formalin-fixed and paraffin-embedded pancreas biopsies were cut into 6 μm consecutive sections and processed for immunohistochemistry for paraffin sections, as previously described (Korsgren et al., 2012). In brief, antigens were unmasked by heat-induced antigen retrieval, using buffer sodium citrate or EDTA according to the manufacturers’ recommendations. Synaptophysin and CD45 or insulin and CD3 double-staining was used for screening for insulitis within the human pancreases. Insulin and CD43 double-staining was used for screening for insulitis within the rat pancreases. Consecutive sections were further stained for CD3, CD4, CD8, CD20, CD68, CD103, insulin and glucagon (table 1). Bound antibodies were visualized using Dako EnVision and diaminobenzidine-based substrate or double stained using EnVision G/2 Double Stain System, Rabbit/Mouse (DAB+/Permanent Red). Sections were counterstained with hematoxylin and analysed by light microscopy Leica. Rat spleen sections were used as positive control for all antibodies. Negative controls had the primary antibody replaced by buffer.

**Table 1.** Detailed list of antibodies used.

| **Name** | **Clone and supplier** | **HIER pH** | **Dilution** |
| --- | --- | --- | --- |
| anti-CD3 (rat) | SP7 (Abcam) | 6.0 | 1:100 |
| anti-CD4 (rat) | CAL4 (Abcam) | 9.0 | 1:4000 |
| anti-CD8 (rat) | OX-8 (Abcam) | 6.0 | 1:1000 |
| anti-CD20 (rat) | SP32 (Abcam) | 6.0 | 1:100 |
| anti-CD43 (rat) | AbD (BioRad) | 6.0 | 1:200 |
| anti-CD68 (rat) | ED1 (Biorad) |  | 1:100 |
| anti-CD103 (rat) | ERP2259027 (Abcam) | 9.0 | 1:1000 |
| anti-Insulin | A0564 (Agilent)) |  | 1:200 |
| anti-Glucagon | LS C312053 (LSBIO) | 6.0 | 1:400 |
| anti-CD45 (human) | 2B11+PD7/26 (Agilent) | 9.0 | 1:75 |
| anti-Synaptophysin | DAK-SYNAP (Agilent) | 9.0 | 1:100 |
| anti-CD3 | Polyclonal (Agilent) | 9.0 | 1:100 |

### Statistical Analyses

Data from the IVGTT are presented as means ± SEM. The statistical significance of the differences between groups was analyzed by the Kruskal-Wallis test followed by Dunn’s test for multiple comparisons. PCR array data were analyzed with non-parametric testing using Qlucore Omics Explorer version 3.3 software with an interface to R (Qlucore, Lund, Sweden). FDR was determined using the Benjamini Hochberg procedure.

## Acknowledgement

The authors are grateful to Karin Fonnaland for excellent technical assistance. Acknowledgement. The study was supported by grants from the Swedish Medical Research Council (2019-01415), Novo Nordisk Foundation, the Ernfors Family Fund, Barndiabetesfonden, Diabetesfonden, the Sten A Olssons Foundation.

## Author Contributions

A.T., S.I., Å.M., O.S. and O.K designed, analyzed and interpreted the study and wrote the manuscript. O.K. is the guarantors of this work and, as such, had full access to all the data in the study and takes responsibility for the integrity of the data and the accuracy of the data analysis.

